# Basal Gp78-dependent mitophagy promotes mitochondrial health and limits mitochondrial ROS

**DOI:** 10.1101/2021.09.17.460825

**Authors:** Parsa Alan, Kurt R. Vandevoorde, Bharat Joshi, Ben Cardoen, Guang Gao, Yahya Mohammadzadeh, Ghassan Hamarneh, Ivan R. Nabi

**Affiliations:** Life Sciences Institute, Department of Cellular and Physiological Sciences, University of British Columbia, Vancouver, BC V6T 1Z3, Canada; School of Computing Science, Simon Fraser University, Burnaby, BC V5A 1S6, Canada

**Keywords:** Gp78 ubiquitin ligase, mitochondria, mitophagy, reactive oxygen species, GFP-mRFP tandem fluorescent-tagged LC3, spot detection, SPECHT

## Abstract

Mitochondria are major sources of cytotoxic reactive oxygen species (ROS) that contribute to cancer progression. Mitophagy, the selective elimination of mitochondria by autophagy, monitors and maintains mitochondrial health and integrity, eliminating ROS-producing mitochondria. However, mechanisms underlying mitophagic control of mitochondrial homeostasis under basal conditions remain poorly understood. Gp78 E3 ubiquitin ligase is an endoplasmic reticulum membrane protein that induces mitochondrial fission and mitophagy of depolarized mitochondria. Here, we report that CRISPR/Cas9 knockout of Gp78 in HT-1080 fibrosarcoma cells increased mitochondrial volume and rendered cells resistant to carbonyl cyanide m-chlorophenyl hydrazone (CCCP)-induced mitophagy. These effects were phenocopied by knockdown of the essential autophagy protein ATG5 in wild-type HT-1080 cells. Use of the mito-Keima mitophagy probe confirmed that Gp78 promoted both basal and damage-induced mitophagy. Application of a spot detection algorithm (SPECHT) to GFP-mRFP tandem fluorescent-tagged LC3 (tfLC3)-positive autophagosomes reported elevated autophagosomal maturation in wild-type HT-1080 cells relative to Gp78 knockout cells, predominantly in proximity to mitochondria. Mitophagy inhibition by either Gp78 knockout or ATG5 knockdown reduced mitochondrial potential and increased mitochondrial ROS. Live cell analysis of tfLC3 in HT-1080 cells showed the preferential association of autophagosomes with mitochondria of reduced potential. Basal Gp78-dependent mitophagic flux is therefore selectively associated with reduced potential mitochondria promoting maintenance of a healthy mitochondrial population and limiting ROS production.

## INTRODUCTION

Mitophagy is a process that selectively targets mitochondria for autophagy and lysosomal degradation^1, 2, 3^. Mitophagy plays a critical role in cellular health by controlling mitochondrial mass and eliminating damaged or dysfunctional mitochondria^4^ Reactive oxygen species (ROS) are generated in mitochondria or by membrane-bound NADPH oxidase and elevated in tumor cells^5, 6^. At high levels, ROS induces DNA mutations leading to cellular transformation and tumorigenesis. Autophagy and, more specifically, mitophagy play key roles in controlling ROS production and thereby cancer progression^7, 8^. Indeed, in response to hypoxia, BNIP3/NIX/FUNDC1-induced mitophagy reduces cellular ROS levels and limits metastasis^2, 9, 10^. However, the relationship between mitophagy and ROS production and its regulation in cancer remains complex and not fully elucidated^11^.

Several ubiquitin-dependent molecular pathways mediate mitophagy, including the well-characterized PINK1/Parkin pathway, closely associated with Parkinson’s disease^12^. In response to mitochondrial damage, including ROS, loss of mitochondrial potential prevents removal of PTEN-induced putative kinase 1 (PINK1) from the outer mitochondrial membrane; PINK1, through phosphorylation, recruits and activates the Parkin E3 ubiquitin ligase to damaged mitochondria, triggering recruitment of the autophagy machinery, including the autophagosome-associated protein, microtubule-associated protein 1A/1B-light chain 3 (LC3/ATG8), and autophagosome formation^13, 14, 15, 16^. Lipidation transforms cytosolic LC3-I to membrane associated LC3-II promoting formation of the phagophore that engulfs dysfunctional mitochondria into autophagosomes for delivery to lysosomes for degradation^17^ However, many cell lines do not express Parkin, tissue expression of Parkin is varied, and mitophagy independent of Parkin and/or PINK1 has been reported^18, 19, 20^. Despite the focus on the role of PINK1 and Parkin in damage-induced mitophagy^21, 22, 23, 24^, both are dispensable for basal mitophagy in various *in vivo* systems^19, 25^. Other molecular mechanisms must therefore regulate basal mitophagy.

Other ubiquitin ligases such as Mul1, MARCH, SMURF, HUWE1, RNF185 and Gp78 have been reported to function independently or in parallel with PINK1/Parkin^26, 27, 28, 29, 30, 31, 32^. PINK1-dependent, Parkin-independent mitophagy pathways include the synphilin/SIAH1 and the Mulan pathways^28, 29, 33^. However, synphilin is primarily expressed in the brain and whether Mulan functions downstream of or independently of PINK1 remains to be determined^1^. Gp78, a key E3 ubiquitin ligase in the endoplasmic reticulum (ER)^34, 35^, exhibits both pro-metastatic and tumor suppressor properties^36, 37^ Upon mitochondrial depolarization, Gp78 degrades mitofusin^30, 38, 39^. Gp78 degradation of outer mitochondrial membrane proteins form mitoplasts that interact with the endoplasmic reticulum and are degraded by reticulophagy^40^. A role for Gp78 in mitophagy is based primarily on loss of mitochondrial mass upon Gp78 overexpression in the presence of the mitochondrial oxidative phosphorylation uncoupler CCCP^30, 38, 39, 40^. Defining the role of Gp78, and other mitophagy effectors, in tumor cell mitophagy requires study of Gp78 knockout cells and of basal, and not damage-induced, mitophagy.

Here, using CRISPR/Cas9 knockout of Gp78 in HT-1080 fibrosarcoma cells, Gp78 is shown to promote both basal and damage-induced mitophagy leading to ATG5-dependent reduction in mitochondrial mass, increased mitochondrial potential and reduced mitochondrial ROS. Application of the spot detection algorithm SPECHT^41^ to monitor flux of the autophagosome reporter tfLC3 localized autophagosome maturation in proximity to mitochondria with reduced membrane potential. By targeting damaged mitochondria for degradation, Gp78-dependent mitophagic flux regulates the homeostasis of healthy mitochondria promoting mitochondrial health and reducing ROS production in cancer cells.

## RESULTS

### Impaired basal and damage-induced mitophagy in Gp78 knockout HT-1080 cells

The HT-1080 fibrosarcoma cell line expresses high levels of Gp78 protein and has been extensively used for the study of Gp78^37, 42, 43^. We previously used stable miRNA and inducible lenti-shRNA approaches to post-transcriptionally knockdown Gp78 mRNA^30, 43, 44, 45^. We now applied CRISPR/Cas9 to knockout Gp78 using two different guided RNA sequences: gRNA1 targets and eliminates the ATG start codon; gRNA2 induces a frameshift sixteen amino acids downstream of the ATG start codon (Figure 1A). Complementary gRNA1 or gRNA2 oligos were annealed and cloned in the GeneArt-OFP plasmid, sequence verified and transfected into HT-1080 cells. We screened sixty clones by Western blotting; from 38 Gp78 knockout clones, three clones showing complete absence of Gp78 from each gRNA were selected (gRNA1: clones #3, 4 and 7; gRNA2: clones #13, 36 and 41). Western blots showed the complete absence of Gp78 in all six gRNA1 and gRNA2 clones compared to wild-type HT-1080 cells (Figure 1A).

**Figure 1:**
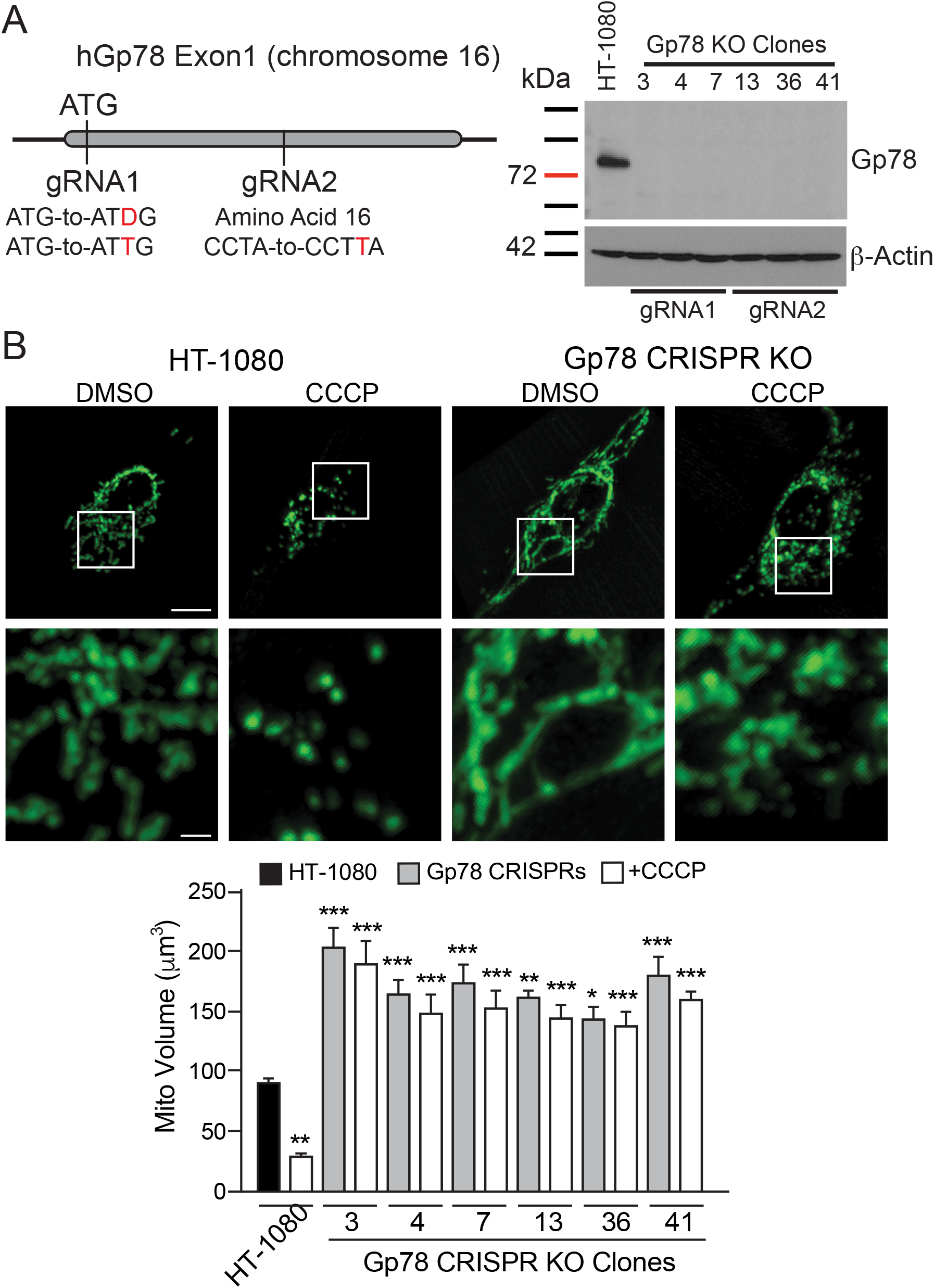
CRISPR/Cas9 knockout of Gp78 in HT-1080 prevents damage-induced mitophagy. (**A**) Schematic showing Exon 1 region of Gp78 gene on chromosome 16 and location of gRNA1 targeting the start codon, deleting ATG (ΔG or inserting extra T), and of gRNA2, inserting an extra T at amino acid 16 causing frameshift and termination. Western blot for Gp78 is shown for wildtype HT-1080 cells and all six gRNA1 and gRNA2 Gp78 knockout CRISPR clones with ß-actin as a loading control. **(B)** Wild-type HT-1080 cells and the six Gp78 knockout CRISPR clones were incubated with either DMSO or 10 μM CCCP for 24 hours then fixed and labeled for mitochondrial ATPB and imaged by 3D spinning disk confocal microscopy. Representative images of HT-1080 and a CRISPR clone including magnification of boxed region are shown. Quantification of total mitochondrial volume is shown in the bar graph (n=3; *, p<0.05; **, p<0.01; ***, p<0.001 relative to DMSO or CCCP treated HT-1080 cells, respectively; ±SEM; Scale Bars: 5 μm; 1 μm for zooms).

Gp78 CRISPR/Cas9 knockout HT-1080 clones labeled for the inner mitochondrial protein ATPB synthase all showed extended mitochondrial networks compared to wild-type HT-1080 cells. Quantification of total mitochondrial volume from 3D spinning disk confocal stacks showed a significant ~2-fold increase in mitochondrial volume in the Gp78 knockout clones relative to wild-type HT-1080 cells (Figure 1B). Rescue of Gp78 knockout HT-1080 cells by transfection of wildtype Gp78, but not Gp78 containing a point mutation in the RING finger domain required for Gp78 ubiquitin ligase activity^34, 46^, reduced mitochondrial volume confirming that Gp78 ubiquitin ligase activity controls mitochondrial volume in HT-1080 cells (Supp. Figure 1). To determine if Gp78 knockout specifically affects mitophagy, we assessed the impact of the mitochondrial membrane potential decoupler CCCP on mitochondrial volume in wild-type and Gp78 knockout HT-1080 cells. CCCP treatment induced mitochondrial fragmentation and a significant reduction in mitochondrial volume in wild-type HT-1080 cells. However, while CCCP treatment induced fragmentation of mitochondria of Gp78 knockout HT-1080 cells, the elevated mitochondrial volume of the Gp78 knockout clones was not reduced (Figure 1B). This suggests that endogenous Gp78 is required for damage-induced mitophagy in HT-1080 cells.

To determine whether autophagy was responsible for the reduced mitochondria levels of Gp78 knockout cells, we knocked down the essential autophagy gene ATG5 in wild-type HT-1080 cells and Gp78 knockout g1-4 and g2-41 clones. By Western blot analysis, mitochondrial ATPB synthase levels showed a significant two-fold elevation in both Gp78 knockout clones relative to wild-type HT-1080 cells (Figure 2A), consistent with the increased mitochondrial volume of these cells (Figure 1B). Upon ATG5 siRNA knockdown, ATPB synthase levels in HT-1080 cells increased to those of Gp78 knockout cells (Figure 2A). Similarly, ATG5 KD significantly increased mitochondrial volume by 3D spinning disk confocal analysis of both untreated and CCCP-treated wild-type HT-1080 cells (Figure 2B). ATG5 KD did not affect mitochondrial volume in Gp78 knockout clones in the absence or presence of CCCP. This suggests that the increased mitochondrial volume of Gp78 knockout cells, in the absence of CCCP, is due to inhibition of active Gp78-dependent mitophagy in wild-type HT-1080 cells.

**Figure 2:**
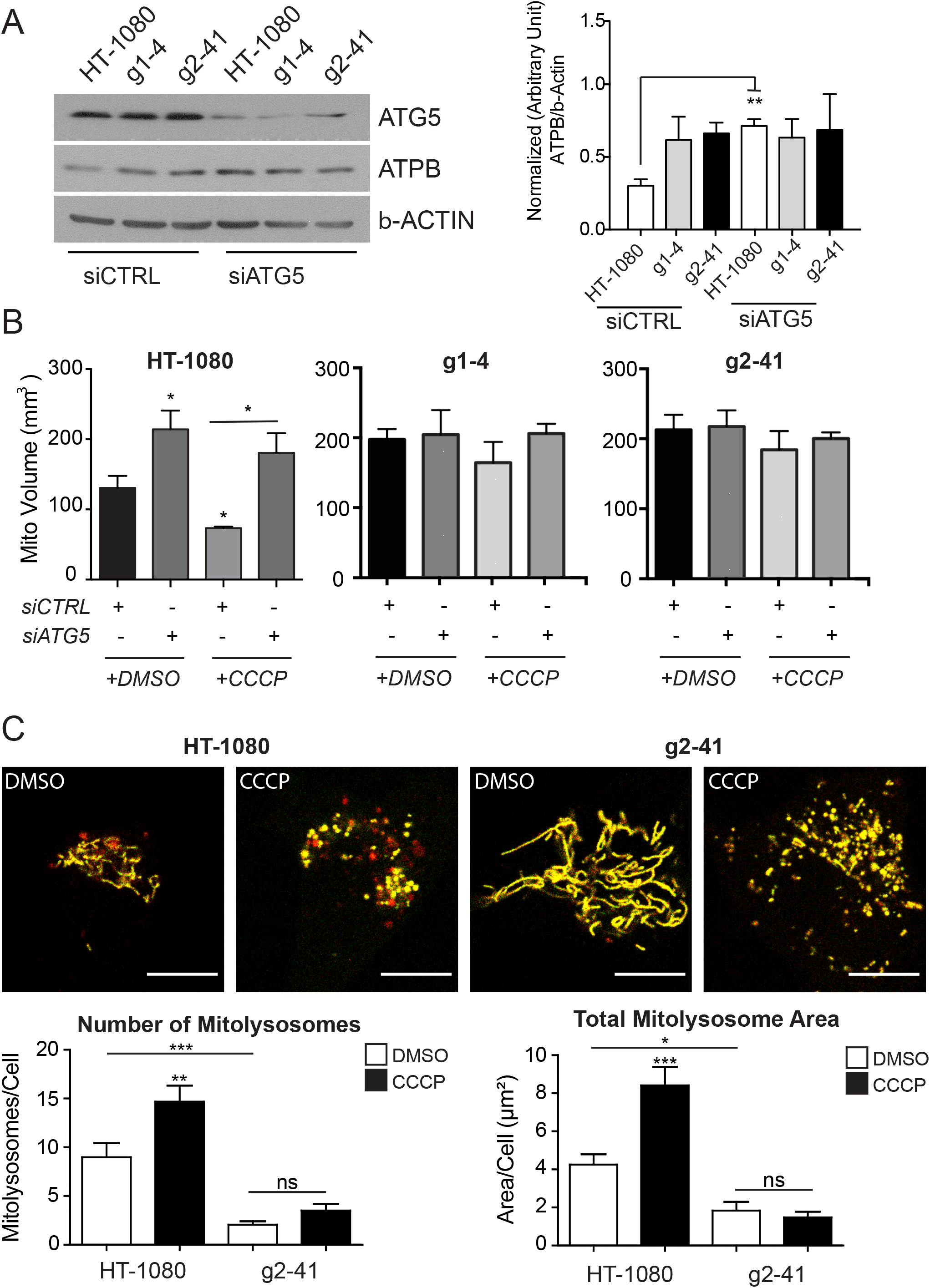
Gp78 induces basal mitophagy in HT-1080 cells. **(A)** Wild-type HT-1080 cells and the g1-4 and g2-41 Gp78 knockout CRISPR clones were transfected with non-specific siCTL or siATG5 and western blotted for ATG5, mitochondrial ATPB and ß-actin. Densitometry quantification of ATBP relative to ß-actin is shown in the bar graph. (t-test, n=3, **p<0.001, ±SEM). **(B)** Wild-type HT-1080 cells and the g1-4 and g2-41 Gp78 knockout CRISPR clones transfected with non-specific siCTL or siATG5 were incubated with either DMSO or 10 μM CCCP for 24 hours and then fixed and labeled for mitochondrial ATPB and imaged by 3D spinning disk confocal microscopy. Quantification of total mitochondrial volume is shown in the bar graph. (n=3, *, p < 0.05; **, p<0.01; ±SEM). **(C)** Wild-type HT-1080 cells and g2-41 knockout CRISPR clones were transfected with mito-Keima and treated with DMSO or CCCP for 24 hours. Confocal fluorescence images show mito-Keima in both neutral (yellow) and low pH (red) environments. Accumulation of mitolysosomes (red) is observed selectively in HT-1080 cells and increased upon CCCP treatment. Quantification of both the number of mitolysosomes per cell and the total area of mitolysosomes per cell is shown in the bar graphs (Scale bar: 10 μm; n=3; *, p<0.05; **, p<0.01; ***, p<0.001; ns, no significance; ±SEM).

We then used the mitoKeima mitophagy probe^47^ to test Gp78 regulation of mitophagy in these cells. Excitation maxima of the Keima fluorescent protein is pH sensitive enabling differential detection of mitochondria-targeted Keima in mitochondria, or following mitophagy and delivery to acidic lysosomes. Live cell images of wild-type and Gp78 knockout HT-1080 cells transiently transfected with mitoKeima were ratio analyzed using an ImageJ plugin^48^ to detect mitochondrial and lysosomal-associated mitoKeima probe. MitoKeima labeled lysosomes are observed in wildtype HT-1080 cells and to a far lesser extent amongst the extended mitochondrial network of Gp78 knockout cells (Figure 2C). CCCP treatment increased the number and area of mitoKeima-positive lysosomes in wild-type HT-1080 cells indicative of damage-induced mitophagy induction. The Gp78 knockout clone showed no increase in mitoKeima lysosome distribution (Figure 2C). Gp78 knockout in HT-1080 cells is therefore associated with impaired basal and damage-induced mitophagy.

### Gp78promotes basal mitophagic flux

We then undertook to determine whether Gp78 regulates autophagic flux and more specifically mitophagic flux. We monitored accumulation of the membrane associated LC3B-II by Western blot upon inhibition of lysosomal acidification and degradation using BafA1 (Figure 3). Cells were treated for 4 hours with CCCP to induce mitophagy or serum-starved to induce macroautophagy. With increasing time of BafA1 incubation, cells treated with DMSO or serum-starved showed increased LC3B-II accumulation in both HT-1080 cells and the g1-4 and g2-41 Gp78 knockout clones. Quantification of flux, based on the slope of LC3B-II band density relative to ß-actin over time, showed no significant differences between the three cell lines at basal levels. Upon treatment with CCCP, LC3B-II flux was observed in HT-1080 cells but not in the Gp78 knockout clones. These data demonstrate that Gp78 is selectively required for damage-induced mitophagic flux, consistent with the inability of CCCP to induce mitochondrial loss in Gp78 knockout cells (Figure 1). However, this Western blot analysis did not detect differences in basal autophagic flux that could explain the increased mitochondrial mass of Gp78 knockout cells, perhaps related to previously reported limitations of Western blot analysis of autophagic proteins for detection of autophagic flux^49, 50^.

**Figure 3:**
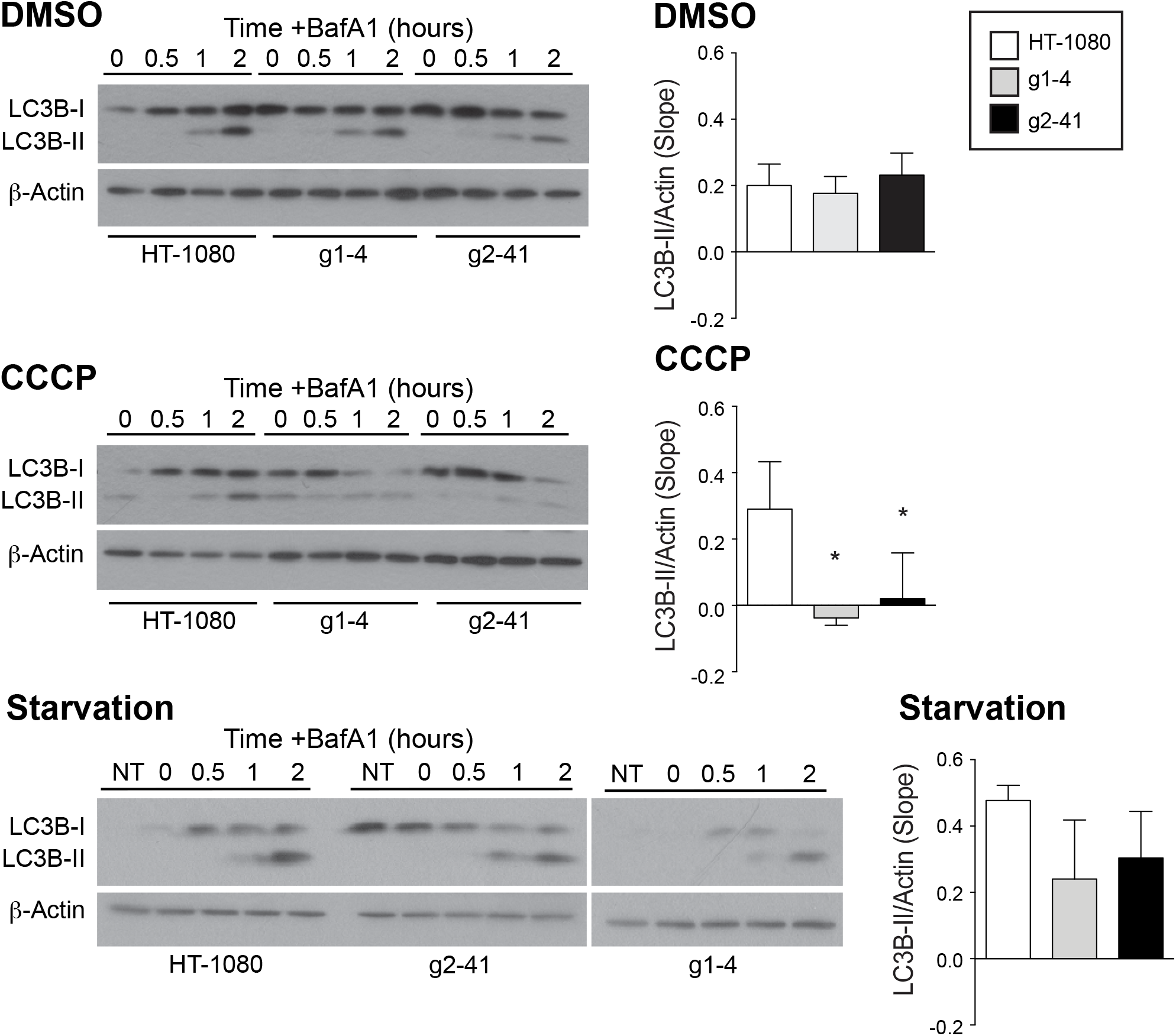
Flux of autophagy protein LC3B-II in wild-type HT-1080 and Gp78 knockout cell lines. Western blot analysis and probing of LC3B-II in HT-1080 cells and g1-4 and g2-41 Gp78 knockout CRISPR/Cas clones after 4 hours treatment with either DMSO, CCCP or starvation in presence of 100 nM of BafA1 for 0, 30, 60 and 120 minutes. DMSO or serum-starved cells show increased and accumulated LC3B-II with BafA1 in time-dependent manner. The slope of LC3B-II band density quantified relative to ß-actin show significant differences between the three cell lines; LC3B-II flux in CCCP treatment of HT-1080 cells but not in the Gp78 knockout clones can be seen (n=4, *, p < 0.05; **; ±SEM).

To develop a more specific and sensitive assay to measure basal autophagic flux, we used the tandem fluorescent-tagged tfLC3 in which LC3 is linked to both GFP and mRFP1; pH-sensitive GFP is a marker for early autophagosomes while mRFP remains fluorescent in acidic autophagolysosomes^51^. Increased dual GFP-mRFP tfLC3 fluorescence upon BafA1 neutralization of acidic autophagolysosomes, enabling GFP fluorescence and preventing GFP degradation, is therefore an indicator of autophagic flux. Stably expressed tfLC3 in HT-1080 cells presented a diffuse cytosolic distribution in both the GFP and mRFP channels with a few puncta corresponding to autophagic vacuoles (Figure 4A). Increased accumulation of mRFP puncta relative to GFP puncta, due to mRFP’s pH-insensitive fluorescence and resistance to lysosomal degradation, is indicative of ongoing basal autophagy in these cells^52, 53^. Dually labeled GFP-mRFP puncta correspond to neutral pH autophagosomes; accumulation of dually labeled GFP-mRFP puncta upon short-term BafA1 treatment reflects the accumulation of intact tfLC3 in acidic autophagosomal compartments and is therefore a measure of autophagic flux. Upon 4-hour CCCP treatment in the presence of BafA1 for the final two hours, conditions that induce a robust mitophagic flux response (Figure 3), an increase in both GFP and mRFP labeled puncta was observed, including multiple overlapping puncta (Figure 4A).

**Figure 4:**
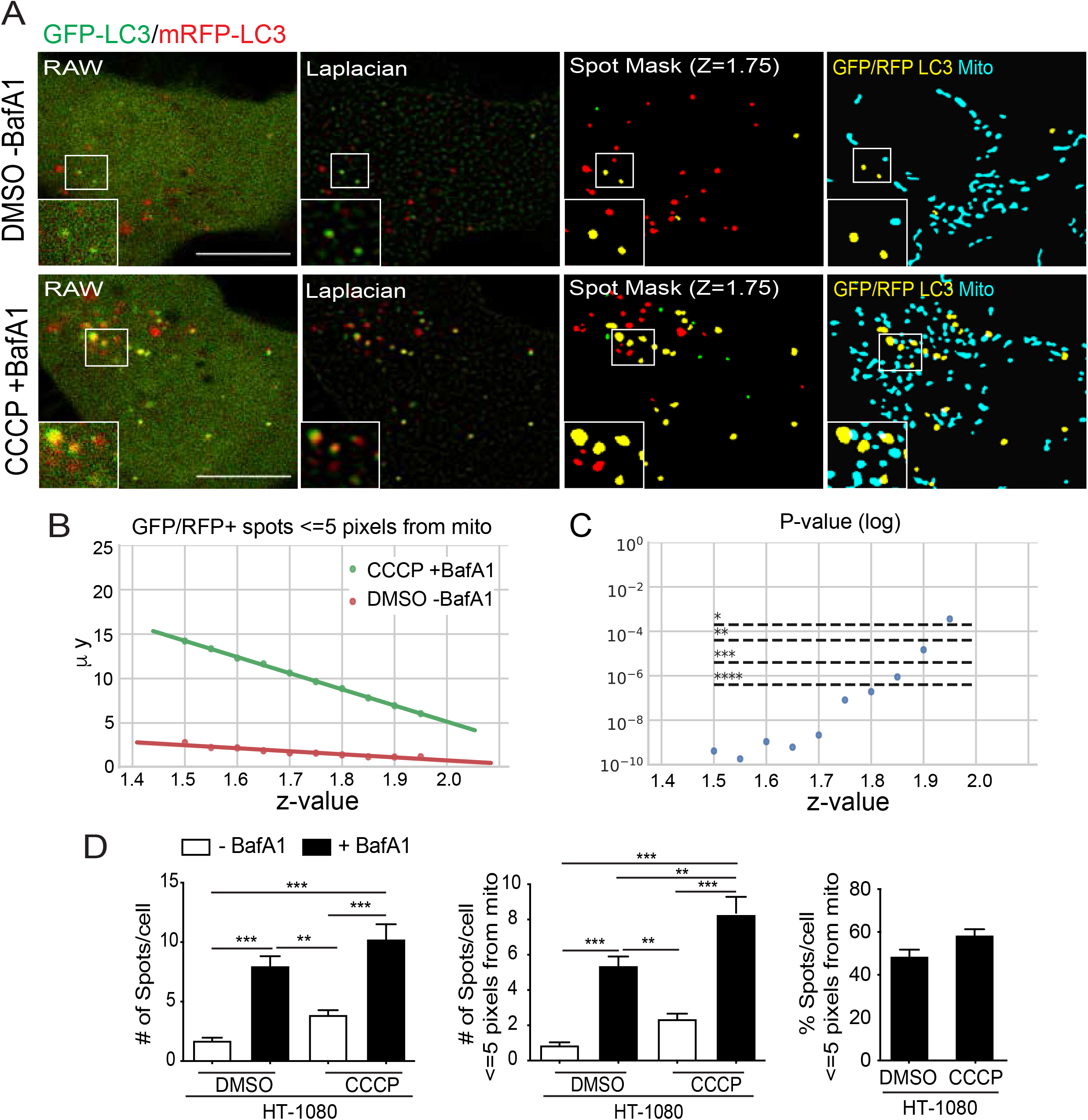
SPECHT spot detection detects LC3 puncta and basal autophagic flux. **(A)** Raw confocal images, Laplacian transforms and SPECHT (z=1.75) spot detection of GFP (green) and RFP (red) tfLC3 signal as well as overlay of GFP-RFP-positive tfLC3 puncta (yellow) with a mask of MitoView mitochondrial labeling (blue) are shown. **(B)** A parameter sensitivity study testing various z-values to determine the significance levels between overlapping GFP-mRFP tfLC3 spots in HT-1080 cells treated with either DMSO or with CCCP and BafA1. Y-axis denotes the mean number of detected spots. **(C)** A parameter sensitivity study testing various z-values and their effect on the mean of number of overlapping GFP-mRFP tfLC3 spots per cell with mitochondria, within or equal to the 5-pixel unit resolution limit. Bonferroni correction with m=252 is applied to correct for multiple hypothesis testing. The Kruskal non-parametric test is applied to test if the samples originate from the same distribution. **(D)** Bar graphs show the number of overlapping GFP-LC3 and mRFP-LC3 spots per HT-1080 cell treated with DMSO or CCCP±BafA1, total and in association with mitochondria, per cell (Mean ± SEM; 10–20 cells/experiment; n = 6, ***p < 0.001, ****p < 0.0001). Scale bar, 10 μm.

However, quantifying LC3 puncta amongst the diffuse fluorescence signal of cytoplasmic LC3 is challenging^54^ Application of the SPECHT spot detection algorithm was used to robustly identify LC3 puncta^41^. To remove puncta smaller than the 250 nm resolution of diffraction limited confocal microscopy that could not correspond to 300-1000 nm autophagosome precursors or phagophores^55^, we applied a size filter that removed any puncta of 25 pixel area (pixel size = 56.6 nm) or less. Overlapping GFP and mRFP puncta (by at least one pixel), considered to be dually-labeled GFP-mRFP puncta (spots), are highlighted in yellow and shown overlaid with the mask of the mitochondrial signal (Figure 5A).

**Figure 5:**
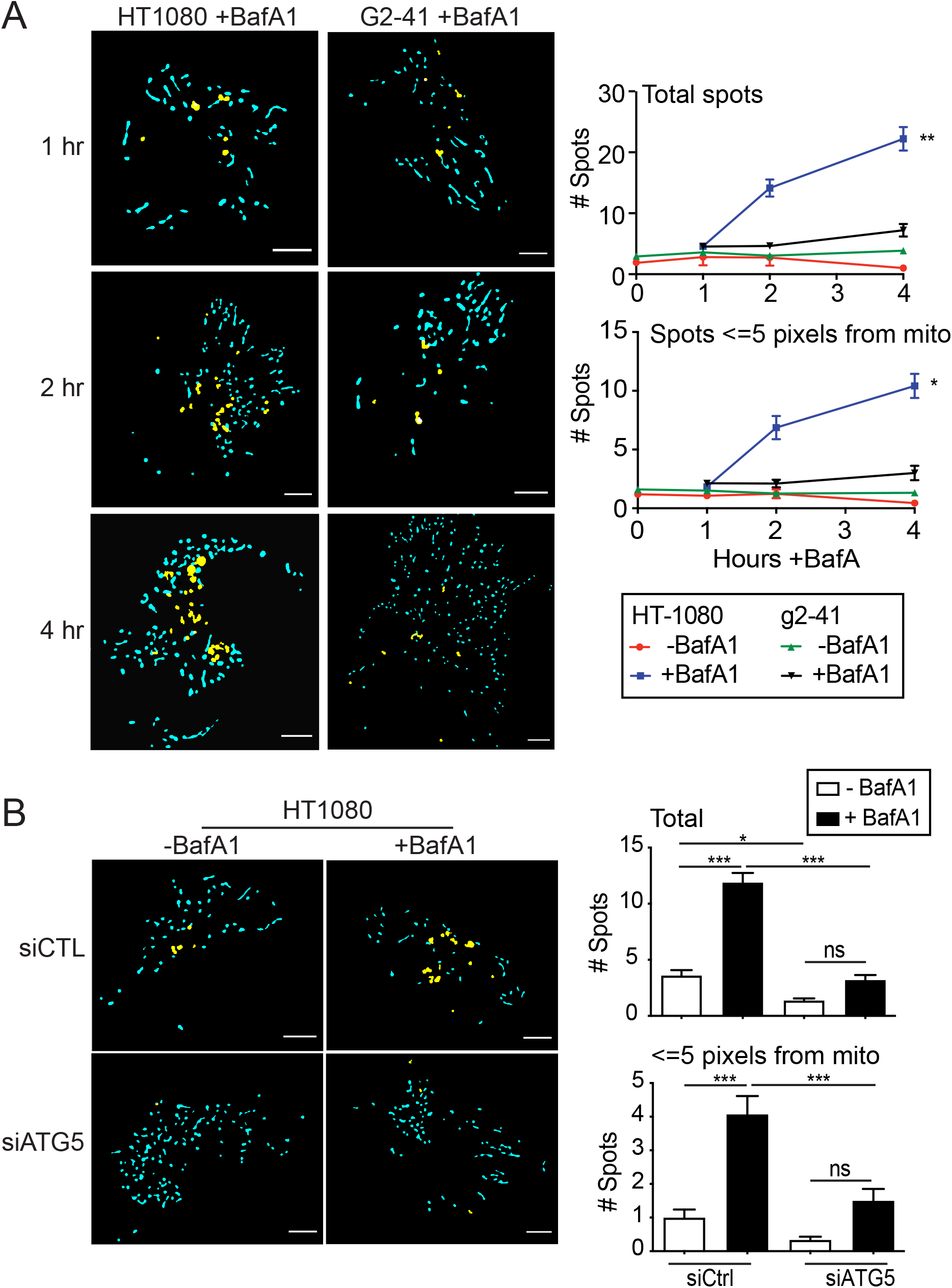
Gp78 knockout in HT-1080 cells results in impaired autophagic flux. **(A)** Masks of GFP-mRFP-positive tfLC3 puncta (yellow) and mitochondria (blue) in live HT-1080 and g2-41 cells stably transfected with tfLC3 and labelled with the mitochondrial dye MitoTracker Deep Red FM. Live cells were either treated with 100 nM BafA1 and imaged at the indicated time points. Bar graphs show the number of GFP-mRFP-positive tfLC3 puncta, total and in association with mitochondria, per cell (Mean ± SEM; n=3, *p<0.05, **p<0.01). Scale bar, 10 μm. **(B)** GFP-mRFP-positive tfLC3 puncta (yellow) and mitochondria (blue) masks of HT-1080 and g2-41 Gp78 knockout cells stably transfected with tfLC3 and labelled with mitochondrial dye MitoTracker Deep Red FM transfected with either control of siATG5 siRNA and treated with 100 nM BafA1. Bar graphs show the number of GFP-mRFP-positive tfLC3 puncta, total and in association with mitochondria, per cell (Mean ± SEM; n=3, *p<0.05, **p<0.01). (Mean ± SEM; n=3, *p<0.05, ***p<0.001). Scale bar, 10 μm.

To identify the intensity-independent z score threshold (see Materials and Methods – spot detection) value that most accurately detects LC3-positive autophagosomes, we performed a parameter sensitivity study on z. We tested the impact of z on differential association of autophagosomes with mitochondria between DMSO and CCCP+BafA1 treated cells, known to present a highly significant difference in autophagic flux (Figure 3). A total of 21 z-values were tested, with values of z > 2 resulting in empty image masks. Given that we have 4 groups (CCCP±BafA, DMSO±BafA), we have 6 possible combinations (from 4 choose 2) and 2 cell lines (m = 6×2×21 = 252 hypothesis tests). To account for multiple testing correction, we applied the Bonferroni correction with a corrected α of α/252 and the Kruskal non-parametric test to compare if the difference between the 2 conditions is significant and consistent across the parameter range. Low z-values causes higher recall of objects, which possibly skews the results towards significance. At the other end, high z-values erode the size and diminishes the counts of detected puncta (Figure 4B, Supp. Figure 2). Application of SPECHT to size filtered LC3 at z=1.75 dramatically improved LC3 puncta detection, significantly detecting differential expression of dually labeled GFP-mRFP tfLC3 puncta between control and CCCP+BafA1 treated cells and accurately reproducing puncta distribution (Figure 4C, Supp. Figure 2). Two hours BafA1 treatment induced dually labeled GFP-mRFP tfLC3 puncta not only in CCCP-treated cells but also in untreated cells, indicative of active basal autophagy in HT-1080 cells. BafA1-induced GFP-mRFP tfLC3 puncta encompass both neutral autophagosomes and acidic autophagolysosomes, that following the sequestration event will move away from mitochondria. About 50% of BafA1-induced GFP-mRFP tfLC3 puncta were located within the resolution limit of 5 pixels (~250 nm) to mitochondria for both untreated and CCCP treated cells (Figure 4D), indicative of a similar mitochondria association of autophagic vacuoles under basal and CCCP conditions. SPECHT analysis of tfLC3 therefore detects both basal and damage-induced mitophagic flux in HT-1080 cells.

Having established conditions to detect mitochondrial-associated autophagosomes in HT-1080 cells in response to CCCP, known to induce mitophagy in HT-1080 cells, we then applied this approach to HT-1080 and Gp78 knockout g2-41 cell lines in the absence of CCCP. Time course analysis of tfLC3 expression showed a significant increase in dual labeled GFP-mRFP LC3 spots upon 4 hours BafA1 treatment of HT-1080 cells in proximity to mitochondria (Figure 5A). Dual labeled GFP-mRFP LC3 spots showed no increase in the absence of BafA1 or in g2-41 Gp78 knockout cells in the presence of BafA1. The BafA1-induced increase in GFP-mRFP-positive tfLC3 spots in the absence of CCCP is indicative of basal autophagic flux and association of tfLC3 puncta to mitochondria suggestive of basal mitophagy. Inhibition of autophagy by siRNA knockdown of the essential autophagy gene ATG5 (siATG5) prevented BafA1 induction of GFP-mRFP-positive tfLC3 puncta, confirming that this assay reports on basal autophagic processes (Figure 5B). Basal mitophagy of HT-1080 cells is therefore Gp78-dependent.

### Gp78 mitophagy promotes mitochondrial health and limits ROS production

We then tested whether Gp78-dependent basal mitophagy impacts mitochondrial potential, and mitochondrial ROS. HT-1080 and g2-41 Gp78 knockout cells were incubated with MitoView 633, a potential-dependent mitochondrial reporter, or MitoSox Red, a mitochondrial ROS reporter and imaged by live cell microscopy. MitoView 633 labeling of HT-1080 cells was significantly reduced by both Gp78 knockout and ATG5 siRNA knockdown (Figure 6A). In contrast, Gp78 knockout and ATG5 siRNA significantly increased MitoSox Red labeling of mitochondrial ROS (Figure 6B). The parallel effect of both Gp78 knockout and ATG5 knockdown indicates that basal mitophagy promotes mitochondrial health and limits ROS production in HT-1080 cells. That siATG5 did not affect either MitoView or MitoSox labeling in Gp78 knockout HT-1080 cells suggests that Gp78 is the key regulator of autophagic mitochondrial quality control in these cancer cells.

**Figure 6:**
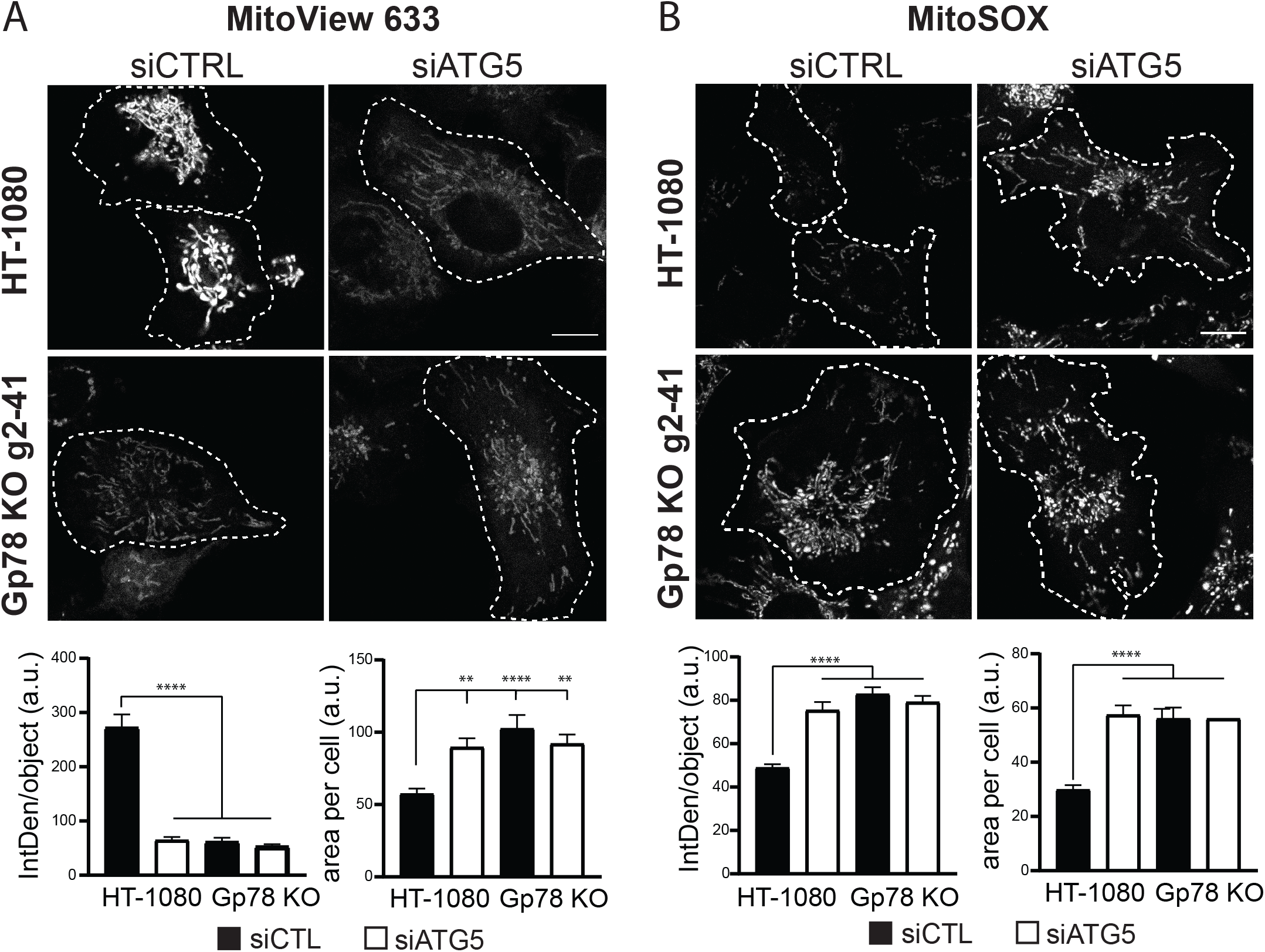
Basal mitophagy maintains healthy mitochondria and limits ROS production in HT-1080 cells. HT-1080 and Gp78 knockout g2-42 HT-1080 cells were transfected with siCTL (black bars) or siATG5 (white bars) and labeled with either the mitochondrial potential reporter MitoView 633 **(A)** or the mitochondrial ROS probe MitoSox **(B)** and (red) for 30 minutes. Live confocal images are shown and MitoView and MitoSox intensity and area quantified per cell. (Scale Bar: 10 μm; n=3;**p<0.01;****p<0.001).

Live cell time lapse imaging of tfLC3-expressing HT-1080 cells labeled for MitoView 633 was performed to determine if autophagosomes specifically associate with damaged mitochondria, i.e. those showing reduced mitochondrial potential. Cells presenting multiple GFP LC3 puncta were imaged by spinning disk confocal and image stacks, 7 images at 500 nm spacing to cover the depth of the cell and minimize overlap of autophagosomes in different Z-sections, were acquired every minute over 40 minutes. SPECHT analysis identified dual-labeled GFP-mRFP tfLC3 puncta, corresponding to phagophores or autophagosomes, in every frame. Representative time lapse images are presented in Figure 7A. Dual-labeled GFP-mRFP tfLC3 puncta were observed to associate with MitoView 633 labeled mitochondria. To determine if autophagosomes were more closely associated with lower potential mitochondria, we quantified the average intensity of MitoView 633 positive pixels overlapping GFP-RFP-positive tfLC3 puncta relative to average MitoView 633 intensity in either the complete mitochondrial segment overlapping the autophagosome or in the whole cell. MitoView-positive pixels overlapping GFP-mRFP-positive tfLC3 puncta presented a below average mitochondrial potential. Time lapse series acquired every 10 seconds show the dynamic association of GFP-mRFP-positive tfLC3 puncta with low potential mitochondria (Figure 7B; Supplemental Video 1). These data show that autophagosomes in HT-1080 cells associate with damaged mitochondria, thereby contributing to maintenance of mitochondrial health of these cells.

**Figure 7:**
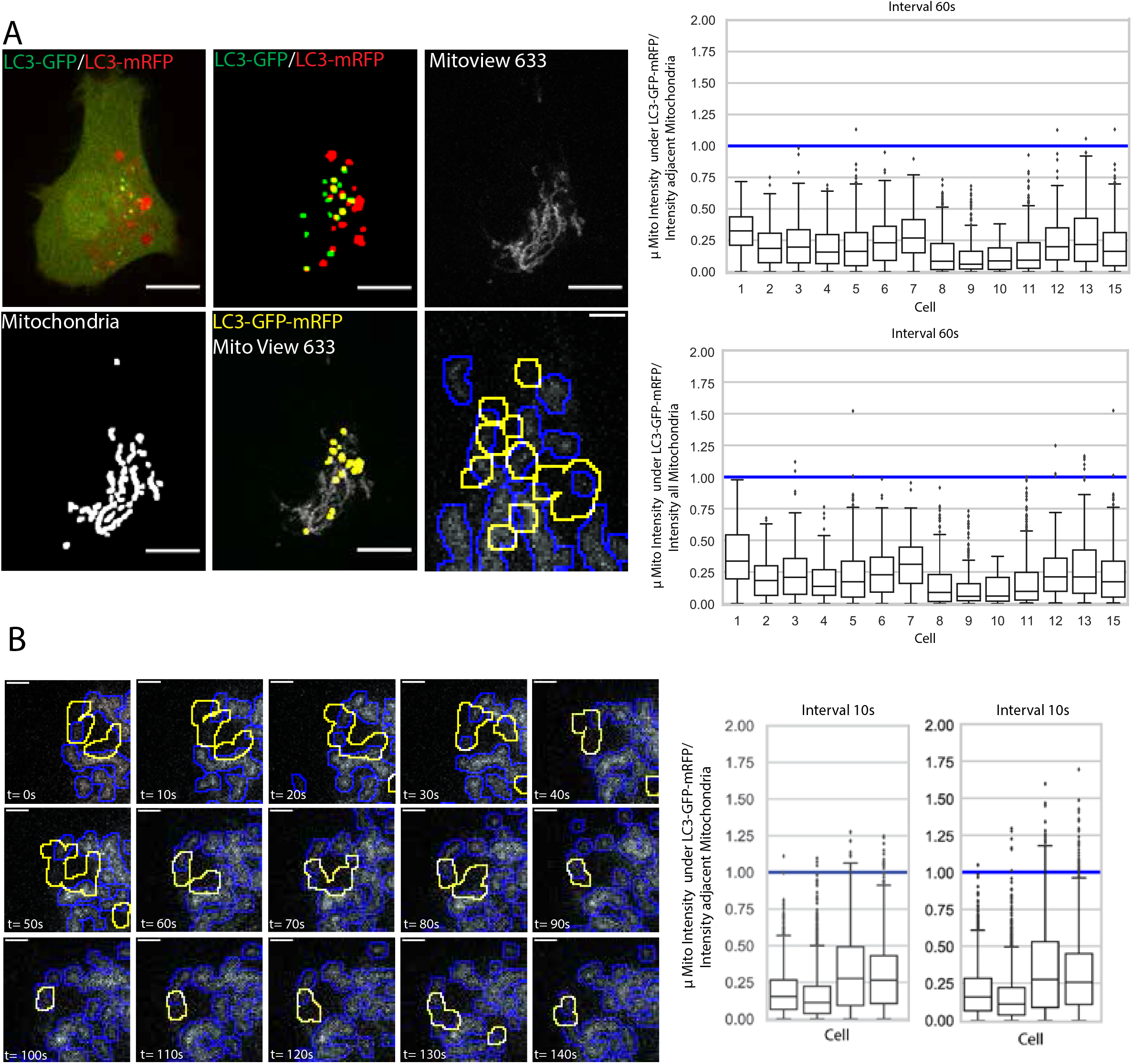
Time lapse imaging of tfLC3 in HT-1080 cells shows autophagosome association with low potential mitochondria. **(A)** Z-stacks (7 images; 500 nm spacing; 14 cells) of HT-1080 cells expressing tfLC3 and labeled with MitoView were acquired by spinning disk confocal every minute over 40 minutes and processed by SPECHT to reveal GFP-mRFP-positive tfLC3 puncta and mitochondria. A representative image of a single Z-section shows SPECHT processing of tfLC3 and mitochondria signals (Scale bar: 10 μm). ROI shows overlap of autophagosomes with lower potential mitochondria (Scale bar: 2 μm). Average intensity of MitoView positive pixels overlapping GFP-mRFP-positive tfLC3 puncta was quantified relative to the average of all MitoView positive pixels in the adjacent mitochondrial segment or in all mitochondrial segments in the cell. For each GFP-mRFP-positive tfLC3 puncta in any of the 7 Z-slices, the ratio of overlapping mitochondria pixels relative to adjacent mitochondria segment and all mitochondria segments in the cell is shown per cell as a boxplot (< 1: fainter than average; > 1 brighter than average). **(B)** Image series from time lapse movies, as described in **(A),** acquired every 10 seconds are shown (see Supplemental video 1). Boxplots are as described in (A). (Scale Bar: 2 μm; n=4).

## DISCUSSION

Here, using CRISPR/Cas9 knockout in the HT-1080 fibrosarcoma cell line, we show that the Gp78 ubiquitin ligase controls basal mitophagic flux and thereby mitochondrial health and ROS production. Basal autophagy maintains cellular homeostasis through the sequestration and subsequent removal of bulk cytoplasmic constituents by delivery to lysosomes for degradation. Functional consequences of defective basal autophagy in ATG5 knockout mice include thyrocyte death, neurodegeneration and increased oxidative stress associated with elevated ROS in these cell populations, possibly because of the accumulation of damaged mitochondria^56, 57^. Basal autophagy has been shown to selectively target subcellular components such as lipids and organelles^58, 59, 60, 61^. The deficient mitochondrial metabolic activity and increased ROS production associated with the impaired basal mitophagy of Gp78 knockout HT-1080 cells provides a definitive link between basal mitophagy, maintenance of mitochondrial health and ROS production by cancer cells.

### Detecting basal mitophagy

As a dynamic, multistep process, autophagic activity is best measured by transit through the path, or autophagic flux^62^; however, measuring basal autophagy, or autophagic flux in the absence of autophagy inducers, is challenging. Mitophagy is monitored through loss of mitochondrial mass; increased mitochondrial volume of HT-1080 cells upon Gp78 knockout and increased mitochondrial mass upon ATG5 siRNA knockdown was indicative of increased basal mitophagy. Reporters like mito-Keima and Mito-Qc report on delivery of mitochondrial proteins to lysosomes and have proven invaluable to monitor mitophagic flux to lysosomes in cells and tissues^19, 47, 63, 64^ Use of mitoKeima and ratio imaging analysis was able to detect the elevated damage-induced and basal mitophagy in wild-type HT-1080 relative to Gp78 knockout cells. However, these approaches report on late stages, i.e. lysosomal delivery and degradation, of mitophagy.

Approaches to measure earlier stages of autophagic flux include analysis of phagophore-and autophagosome-associated LC3B-II and targeted knockdown or knockout of essential autophagy proteins^54^ Western blotting of expression levels of autophagy proteins, such as LC3B-II, report on autophagic activity in a cell population but are limited in their ability to directly measure flux^50^. Western blotting of LC3-II in BafA treated cells detected autophagic flux in response to serum starvation and, selectively for wild-type but not Gp78 knockout cells, upon CCCP treatment. This supports a selective deficiency in mitophagy and not general autophagy upon Gp78 knockout. However, Western blotting for LC3B-II was not sensitive enough to detect differences in basal autophagic flux between cell types.

Fluorescent LC3 expression reporters such as GFP-LC3 and tfLC3 have been used to study autophagic flux in cells and tissues^19, 47, 63, 64^ Tandem tfLC3 reports on autophagic flux at the level of autophagosome maturation due to the differential sensitivity of GFP and mRFP to acid quenching of fluorescence and degradation in lysosomes. However, analysis and interpretation of the acquired data remain challenging. Segmenting LC3 puncta from cytoplasmic LC3 fluorescence is difficult, such that, in some cases, manual counting of LC3 puncta is required {Klionsky, 2016 #6742^;^ Musiwaro, 2013 #6764}. By applying the spot detection stage of the SPECHT algorithm {Cardoen, 2021 #6763}, we report the sensitive and robust detection of tfLC3 puncta. Using BafA1 inhibition of autophagosome maturation, we demonstrate the presence of basal autophagic flux in HT-1080 cells but not in Gp78 knockout HT-1080 cells. Mitochondrial association of BafA1-induced autophagosomes paired with reduced mitochondria volume and potential and increased mitochondrial ROS upon inhibition of autophagy using siATG5 demonstrates that HT-1080 cells undergo active basal mitophagy. Live cell time lapse analysis of tfLC3 in HT-1080 cells allowed us to demonstrate that autophagosomes associate with reduced potential mitochondria, confirming a role for mitophagy in mitochondrial quality control. The enhanced sensitivity of detection of tandem tfLC3 using SPECHT^41^, even when channels differ markedly in signal to noise ratio, renders this a valuable tool to study basal autophagy in various cell and tissue systems.

### Gp78, mitophagy and ROS production

Gp78 overexpression recruits LC3 to mitochondria-associated ER^30^ and Gp78 has been shown to selectively regulate rough ER-mitochondria contacts^43^. Also, Gp78 has been recently shown to be involved in ERphagy, and selective degradation of the outer mitochondrial membrane, forming mitoplasts and triggering mitophagy^67^. The relationship between Gp78-dependent mitoplast formation, the ER and the dynamic movement and maturation of LC3-positive autophagosomes in proximity to mitochondria that we report here remains to be determined. Considering the dispensable role of Parkin and PINK1 in basal mitophagy *in vivo*^19, 25^, further analysis of the role of Gp78-dependent basal mitophagy in tissue development and cancer progression is warranted.

ROS generation shows a complex association with tumor progression. ROS are critical for tumor initiation and progression acting through activation of oncogenic signaling and DNA mutation; at the same time, increasing ROS production eliminates cancer cells by inducing cell death programs^68^. Regulation of mitochondrial ROS production by BNIP3, a hypoxia-induced mitophagy regulator, suppresses breast cancer progression^69, 70^. The formation of spontaneous liver tumors after 12 months in Gp78 knockout mice supports a tumor suppressor function for Gp78^36^ that is consistent with Gp78-dependent mitophagic control of ROS production. However, Gp78 knockdown in the HT-1080 cells studied here did not affect tumor growth but did inhibit metastasis^37^; further, impaired metastasis was rescued by wild-type Gp78 but not a RING finger Gp78 mutant with deficient ubiquitin ligase activity that does not induce mitophagy^30, 37^ Indeed, Gp78 was shown to promote metastasis though targeted degradation of the metastasis suppressor KAI1^37^ and Gp78 downregulation of KAI1 shown to be associated with mammary gland hyperplasia^71^. This suggests multiple roles for Gp78 in cancer progression, including but not limited to regulation of ROS production through mitophagy.

## MATERIALS AND METHODS

### Antibodies and reagents

Anti-ATPB pAb/mAb (ab128743/ab5432) was purchased from Abcam (Cambridge, MA, USA), anti-ATG5 pAb (NB110-53818) from Novus Biologicals (USA), anti-Gp78 pAb (16675-1-AP) from Proteintech (USA) and anti-LC3B pAb (2775S) from Cell Signaling. Anti-ß-Actin mAb (A5441), tissue culture grade DMSO (D2650), Bafilomycin A1 (Cat# C1988) and CCCP (C2759) were purchased from Sigma-Aldrich (USA). mRFP-GFP tandem fluorescent-tagged LC3 (tfLC3) was a gift from Tamotsu Yoshimori (Addgene plasmid # 21074; www.addgene.org/21074)^51^. mKeima-Red-Mito-7 plasmid was a gift from Michael Davidson (Addgene plasmid #56018; www.addgene.org/56018). MitoTracker Deep Red FM, Live Cell Imaging Solution (A14291DJ), Glucose Solution (A2494001), MitoSOX Red (M36008) and MEM Non-Essential Amino Acids Solution (11140050) were purchased from ThermoFisher (USA). MitoView 633 (#70055) was purchased from Biotium (USA).

### Cell lines and CRISPR/Cas9 knockout of Gp78

The HT-1080 fibrosarcoma cell line was acquired from ATCC, authenticated by Short Tandem Repeat (STR) profiling at the TCAG Genetic Analysis Facility (Hospital for Sick Kids, Toronto, ON, Canada www.tcag.ca/facilities/geneticAnalysis.html), tested regularly for mycoplasma infection by PCR (ABM, Richmond, BC, Canada) and maintained in RPMI 1604 media supplemented with 10% FBS and 1% L-Glutamine in a 37°C incubator with 5% CO2.

GeneArt-CRISPR/Cas9 Nuclease vector with OFP (Orange Fluorescence Protein) kit (A21174) was from Life Technologies (Invitrogen, USA). We used http://crispr.mit.edu to design guided RNAs and http://www.rgenome.net/cas-offinder/ (RGEN tools) to check them for off-target effects and used the following oligonucleotides for guide RNA1 (5’-CAC CGG AGG AAG AGC AGC GGC ATG G-3’, 5’-AAA CCC ATG CCG CTG CTC TTC CTC C-3’) and guide RNA2 (5’-CAC CGG CCC AGC CTC CGC ACC TAC A-3’, 5’-AAA CTG TAG GTG CGG AGG CTG GGC C-3’). From isolated genomic DNA for each clone, the DNA fragment flanking Exon1 of Gp78 was PCR amplified, cloned and sequenced. For gRNA1 clones #3 and #4 the G was deleted from the ATG start codon while for clone #7 an additional T was inserted in the start codon generating ATTG. For all three gRNA2 clones (#13, 36, 41), a T was inserted at amino acid 16 (CCTA to CCTTA) causing a frameshift mutation. The guide RNAs were in vitro annealed, cloned into the GeneArt linear vector according to the supplier’s protocol and sequence verified prior to transfection into HT-1080 cells. Sequence verified gRNA1 or gRNA2 containing GeneArt-CRISPR/Cas9 Nuclease vector with OFP were transiently transfected into HT-1080 cells, plated 24 hours previously, using Effectene transfection reagent (301425, Qiagen, USA). After 36 hours incubation, cells were harvested and genomic DNA isolated to perform GeneArt Genomic Cleavage Detection assay (A24372, Invitrogen, USA) to check cleavage efficiency. Once cleavage efficiency was confirmed, HT-1080 cells were replated for 36 hours, trypsinized, FACS sorted and OFP expressing cells were singly plated in 96 well pates by serial dilution. Single colonies were replicated in 12 well plates; one set was frozen and stored in liqN2 and the other set subjected to lysate preparation, SDS-PAGE and Gp78 western blot analysis. Arbitrarily chosen representative clones (g1-3, g1-4, g1-7; g2-13, g2-36, g2-41) from both gRNAs were expanded, tested for mycoplasma and stored as multiple freeze-downs. From isolated genomic DNA, an approximate 800bp fragment flanking Exon1 of Gp78 was PCR amplified using Q5 (Qiagen, USA) the following primer set (Forward: 5’-CTG GAG GCT ACT AGC AAA-3’, Reverse: 5’-ATG TGG CCC AGT ACC T-3’) and TA cloned. At least ten clones were sequenced from each to confirm INDEL.

HT-1080 and Gp78 CRISPR/Cas9 knockout clones were grown only up to six passages. Cells were passed every 48 hours at a density of 300,000 cells per 10 cm petri dish, rinsed every 24 hours with 10 ml of PBS and supplied with 10 ml of fresh complete medium. HT-1080 cells and the g2-41 Gp78 CRISPR/Cas9 knockout clones were stably transfected with mRFP-GFP tandem fluorescent-tagged LC3 (tfLC3) plasmid using Effectene (Cat. #301425, Qiagen, USA) following the manufacturer’s protocol. After 24 hours incubation, transfected cells were selected against G418 (400ug/ml) for about 14 days. The resistant cell population was pooled and maintained in 50 μg/ml G418.

### siRNA knockdown, plasmid transfection and western blotting

siControl and siATG5 (Cat# D-001810-01-05, Cat# L-004374-00-0005) were purchased from Dharmacon and transiently transfected wherever indicated to wild-type HT-1080 cells or g1-4 or g2-41 Gp78 CRISPR/Cas9 knockout clones using Lipofectamine 2000 (Cat# 11668019, Invitrogen, USA) following the manufacturer’s protocol. All siRNA transfection experiments were for 48 hours and treatments were performed 24 hours post siRNA transfection. Alternatively, cells were transiently transfected with mammalian protein expressing plasmids using Effectene (Qiagen, Germany) following the manufacturer’s protocol. Where indicated, cells were treated with 10 μM of CCCP or a corresponding volume of DMSO as control 24 hours prior to fixation or harvesting cells. Western blotting was performed as previously described using Horseradish Peroxidase (HRP)-conjugated secondary antibody followed by addition of ECL (GE Healthcare Bio-Sciences Corp., USA) to reveal chemiluminescence^71^. Densitometry quantification was done using ImageJ (https://imagej.nih.gov/ij/docs/faqs.html#cite) software.

### Fluorescent labeling of mitochondria

For immunofluorescent labeling, cells were: 1) fixed with 3.0% PFA for 15 minutes at room temperature and washed with PBS-CM (phosphate buffer solution supplemented with 1 mM CaCl2 and 10 mM MgCl2); 2) permeabilized with 0.2% Triton X-100 for 5 minutes and washed with PBS-CM; 3) blocked with 1% BSA for 1 hour at room temperature; 4) labeled with anti-ATPB for one hour followed by washing with PBS-CM; 5) incubated with secondary antibodies for 1 hour followed by washing with PBS-CM; and 6) mounted in ProLong Diamond (ThermoFisher) and cured for 24 hours at room temperature before imaging. Confocal image stacks were obtained on a III-Zeiss spinning disk confocal microscope with either Zeiss Plan-Apochromat 63X/1.2NA or 100X/1.4NA oil objectives using SlideBook 6.0 image acquisition and analysis software (Intelligent Imaging Innovation Inc). ATPB label was thresholded from 3D images to measure mitochondrial volume with SlideBook 6.0 image analysis software.

To assess the impact of Gp78-dependent basal mitophagy on mitochondrial health and mitochondrial ROS, wildtype HT-1080 and the g2-41 Gp78 CRISPR/Cas9 knockout clone were plated into an Ibidi chamber for 24 hours. Cells were then transiently transfected with siRNA targeting ATG5 for 48 hours and labelled with either mitochondrial health sensor dye MitoView633 or mitochondrial ROS dye MitoSOX, at concentrations of 50 nM and 2.5 μM, respectively, for half an hour, washed 3X with warm PBS and incubated in Molecular Probes Live Cell Imaging Solution. Live-cell imaging was performed at 37°C with a Leica TCS SP8 confocal microscope with a 100×/1.40 Oil HC PL APO CS2 objective (Leica, Wetzlar, Germany) equipped with a white light laser, HyD detectors, environmental chamber and Leica Application Suite X (LAS X) software. Images were analyzed using ImageJ software to identify integrated densities of mitochondrial objects as well as the total area of the mitochondrial label, per cell.

### Mitophagic flux assays

In order to study mitophagic flux in HT-1080 or Gp78 CRISPR/Cas9 knockout (g1-4, g2-41) cells, early passage cells (420,000 cells per well) were plated in six well plates for 20 hours, then washed with 1X PBS and treated with DMSO or CCCP in regular medium or medium lacking serum for 4 hours. For each treatment, cells were challenged with 100 nM of BafA1 (Sigma) for 0, 30, 60 or 120 minutes prior to the end of the 4-hour incubation period. Incubation was stopped by washing cells with ice-cold 1X PBS; cells were then harvested on ice lysed with M2-Lysis buffer^72^ supplemented with phosphatase and protease inhibitors tablets (Roche), and lysates ran on 15% SDS-PAGE at constant voltage (75V for fifteen minutes followed by 90V for 2 hours). Separated proteins were electrotransferred onto 0.2 μ pore size PVDF membrane (BioRad), fixed with 0.1% glutaraldehyde in PBST (0.2%) for 30 minutes, blocked with 5% milk in PBST and immunoprobed for LC3B-I and II and ß-Actin. LC3B-II and ß-actin bands were densitometrical quantified using ImageJ software, normalized and statistically analyzed.

To monitor mitophagic flux with mito-Keima, HT-1080 and Gp78 CRISPR/Cas9 knockout g2-41 cells (8,000 cells per well) were plated in an ibidi chamber for 24 hours and then transfected with mito-Keima plasmid^47^ using Effectene transfection reagent (301425, Qiagen, USA). After 24 hours, cells were washed with PBS and treated with DMSO or CCCP in regular medium. Following a 24-hour incubation, the cells were washed 3X with PBS and then incubated in Live Cell Imaging Solution supplemented with 10% FBS, L-glutamine, D-glucose, and MEM Non-Essential Amino Acids Solution prior to imaging on a Leica TCS SP8 confocal microscope equipped with a 100x/1.40 Oil HC PL APO CS2 objective (Leica, Wetzlar, Germany), white light laser and HyD detectors (Leica, Wetzlar, Germany). mito-Keima in a neutral pH environment was detected by excitation at 470nm and in an acidic environment by excitation at 561nm. The HyD detector was open from 592nm to 740nm and equipped with a time gate limiting detection from 0.3ns to 6.5ns after laser activation. To quantify the presence of mitolysosomes, the fluorescent images were loaded into FIJI with the mito-QC Counter macro installed^48^. Settings (Radius for smoothing = 2.5, Ratio threshold = 1.6, Red channel threshold = 3.2) for ratio analysis were determined to report most accurately on mitolysosome expression across all data sets. For some images, large numbers of spots were detected that did not correspond to observed mitolysosomes. For consistency, the two images presenting the largest number of mitolysosomes per group were removed from the analysis.

To monitor autophagic flux with tfLC3, stably transfected HT-1080 and Gp78 CRISPR/Cas9 knockout g2-41 cells were plated overnight and then treated with either DMSO or CCCP for 4 hours and with or without 100 nM BafA1 for the final 2 hours of the incubation period. Mitochondria were labelled with MitoTracker Deep Red FM half an hour prior to the end of the total incubation period. After incubation, cells were gently washed 3X with warm PBS and then incubated in warm Live Cell Imaging Solution just prior to image acquisition. Live-cell imaging was performed using Leica TCS SP8 confocal microscope with a 100×/1.40 Oil HC PL APO CS2 objective (Leica, Wetzlar, Germany) equipped with a white light laser, HyD detectors, and Leica Application Suite X (LAS X) software. Image acquisition was performed in a temperature-controlled system set to 37°C.

For time lapse imaging, HT-1080 cells expressing tfLC3 were plated in ibidi chambers in Molecular Probes Live Cell Imaging Solution supplemented with 10% FBS, L-glutamine, D-glucose, and MEM Non-Essential Amino Acids Solution and labeled with MitoView prior to imaging at 37°C with the 100X/NA 1.45 PL APO objective (Zeiss) of a 3i Yokogawa X1 spinning disk confocal. Image stacks of 7 images with a 500 nm Z spacing were acquired every minute for 40 minutes with a QuantEM 512SC Photometrics camera. Average intensity of MitoView positive pixels overlapping each GFP-mRFP-positive tfLC3 puncta was assessed relative to average intensity of all MitoView-positive pixels in either the adjacent segmented mitochondria or in the cell.

### tfLC3 spot detection analysis (SPECHT)

To identify tfLC3 labeled autophagic vacuoles (autophagosomes), we applied the SPECHT object detection method, that is consistent across channels and robust to intensity variations^41^. SPECHT evolved from the ERGO software for density detection in single molecule localization microscopy^73^ and accepts as input a confocal image and produces, for each channel, a binary mask of detected fluorescent marker concentrations (spots or puncta). SPECHT leverages the Laplacian-of-Gaussian (LoG) object detection method, but ensures detection is adaptive to the image intensity distribution by computing an automatic threshold to postprocess LoG detected objects. The user can express a preference for recall or precision, which SPECHT then translates into channel/image specific threshold values. This preference is referred to in this manuscript as ‘z-value’. A higher value increases precision, at the cost of recall. A lower value can result in higher recall, at cost of precision. To ensure no artificial objects are introduced, the isotropic Gaussian std. dev. was set to round(precision/2) = 3 pixels (pixel size = 56.6 nm). Objects with area smaller than 25 pixels were removed to avoid false counting of artifacts below the precision limit of the acquisition. Distances between objects were measured using Euclidean distance (pixels) between the closest edges of nearest objects. Puncta within 5 pixels (ceil(precision)) of mitochondria (i.e. the resolution limit of ~250 nm) cannot be distinguished from overlapping puncta and were thus counted as overlapping. The area of objects is represented by pixel mask counting. When a red mRFP and green GFP object (puncta) shared a non-zero intersection, the union of the red-green overlapping puncta were considered to correspond to early, neutral pH autophagosomes. Colocalized GFP-mRFP tfLC3 puncta upon BafA1 treatment encompass acidic autophagolysosomes and an increase in GFP-mRFP tfLC3 puncta following BafA treatment is a measure of autophagic flux. We also quantified the number of BafA1-induced GFP-mRFP overlapping tfLC3 puncta within 5 pixels of mitochondria.

To ensure single cell analysis, ROIs encompassing complete, individual cells within the field of view were manually segmented (Figure 6). While this was feasible for the single time point analysis, for the time lapse series (Figure 7), cells were moving and we therefore added an automated preprocessing stage to obtain cell segmentation masks. Input to SPECHT was a sequence of 2D images (1 Z-slice), 3 channels per timepoint. We recover the outline by applying a median filter (window sizes 3×3, 5×5, 9×9) after filtering out the 90% intensity distribution quantile, binarizing the resulting image, and detecting disjointed objects separated by black (filtered) background using the connected components algorithm, such that the sole complete cell will be the largest object. To accommodate the highly fluctuating intensity distribution of live cell imaging over time, we enable SPECHT’s autotuning mode configured to recover all possible objects (recall/precision ratio = 3.75). To prevent inclusion of false positives we: 1) compute the effect size (Cohen’s d) of its intensity distribution with respect to that of the cell and discard any object with a negative effect size; 2) we use the heuristic that the local maxima contained within each detected object, should be a statistical outlier with respect to the overall intensity distribution (Q3 + 1.5 IQR, respectively 3^rd^ quartile and interquartile range) and discard objects that have a maximum intensity that is not at the extremum of the intensity distribution; 3) objects of area smaller or equal to 4 pixels are discarded, as they cannot be shown to be observable under the precision of the system (2 pixels). To ensure we do not compromise objects at the cell edge we widen the cell mask by a dilation operation 4 times (2 x system precision of 2 pixels). A closing operation ensures no holes are left in the cell mask should one channel have no or weak labelling in part of the resulting mask. To ensure our segmentation is valid, without the user having to screen each image, we test that the cell mask is consistent across channels. In addition, we disregard processing of any image where the cell mask touches the border of the image. The combination of high recall followed by high precision filtering, ensures a balanced, robust automatic pipeline for object detection designed for the live cell imaging data. For each GFP-mRFP overlapping spot (C12), we compute the mean mitochondria intensity it overlaps relative to the mean intensity of the associated mitochondria segment as well as the mean intensity of all mitochondria segments in each given 2D image (C). We then compute a box plot of ratios 1 and 2 for all C12 objects, for each cell (Figure 7), where each cell is represented by 7 2D images (per Z-slice), over 40 timepoints. Output is saved in csv files for statistical analysis and postprocessing. The processing code is being prepared for open-source release (Affero GPLv3), and available upon reasonable request.

### Statistical analyses

One-way ANOVA with Dunnett’s multiple comparison test was used for both the fixed and livecell ROS experiments. One-way ANOVA with Tukey’s multiple comparison test was used for the live cell tfLC3 flux, the mitoKeima experiments, the spinning disk mitochondrial volume experiments and the Western blot flux experiments. A 2 tailed t-test was applied for the siATG5 blots. Statistical analyses were performed using GraphPad Prism 6.0 software.

## ACKNOWLEDGEMENTS

This study was supported by a grant from the Canadian Institutes of Health Research (CIHR Grant PJT-148698) and UBC Fellowships (GG). Imaging was performed in the *LSI IMAGING* facility of the Life Sciences Institute of the University of British Columbia using infrastructure funded by the Canadian Foundation of Innovation and BC Knowledge Development Fund as well as a Strategic Investment Fund (Faculty of Medicine, University of British Columbia). This research was enabled in part by support provided by Westgrid (https://www.westgrid.ca/) and Compute Canada (www.computecanada.ca).

