## Supplemental Material for "Basal Gp78-dependent mitophagy promotes mitochondrial health and limits mitochondrial ROS"

### SUPPLEMENTAL FIGURE AND VIDEO LEGENDS

**Supplemental Figure 1: Gp78 rescues mitochondrial phenotype of Gp78 KO HT-1080 cells in a RING domain-dependent manner. (A)** Representative maximum projections of mitochondria labelled with TOMM20 from wild type HT-1080 and Gp78 KO clones transiently transfected with wild type FLAG-Gp78, catalytically inactive RING finger (RF) mutant Gp78, or empty vector pcDNA3. **(B)** Bar graph shows quantification of total mitochondrial volume per cell (n=3; *, $p<0.05$; **, $p<0.01$; ***, $p<0.001$, ****, $p<0.0001$; Mean ± SEM; Scale bar: 10 µm).

**Supplemental Figure 2: The effect of variable Z-values on SPECHT spot detection.** Raw images of grey scale and merged GFP-mRFP tfLC3 of DMSO and CCCP/BafA1 treated HT-1080 cells. Corresponding Laplacian filter is presented with its resultant z-values of 1.50, 1.75, 2.0. Scale bar, 10 µm.

**Supplemental Figure 3:** Full Western blots for the Gp78 and β-actin blots included in Figure 1A.

**Supplemental Figure 4:** Full Western blots for the ATG5, ATPB and β-actin blots included in Figure 2A.

**Supplemental Figure 5:** Full Western blots for the LC3-B and β-actin blots for DMSO, CCCP and starvation treated cells included in Figure 3.

**Supplemental Video 1**: Time lapse movie of GFP-mRFP-positive tfLC3 puncta (yellow) and MitoView labeled mitochondria in HT-1080 cell (blue SPECHT outline overlaid on raw MitoView image). Image acquired every 10 seconds.

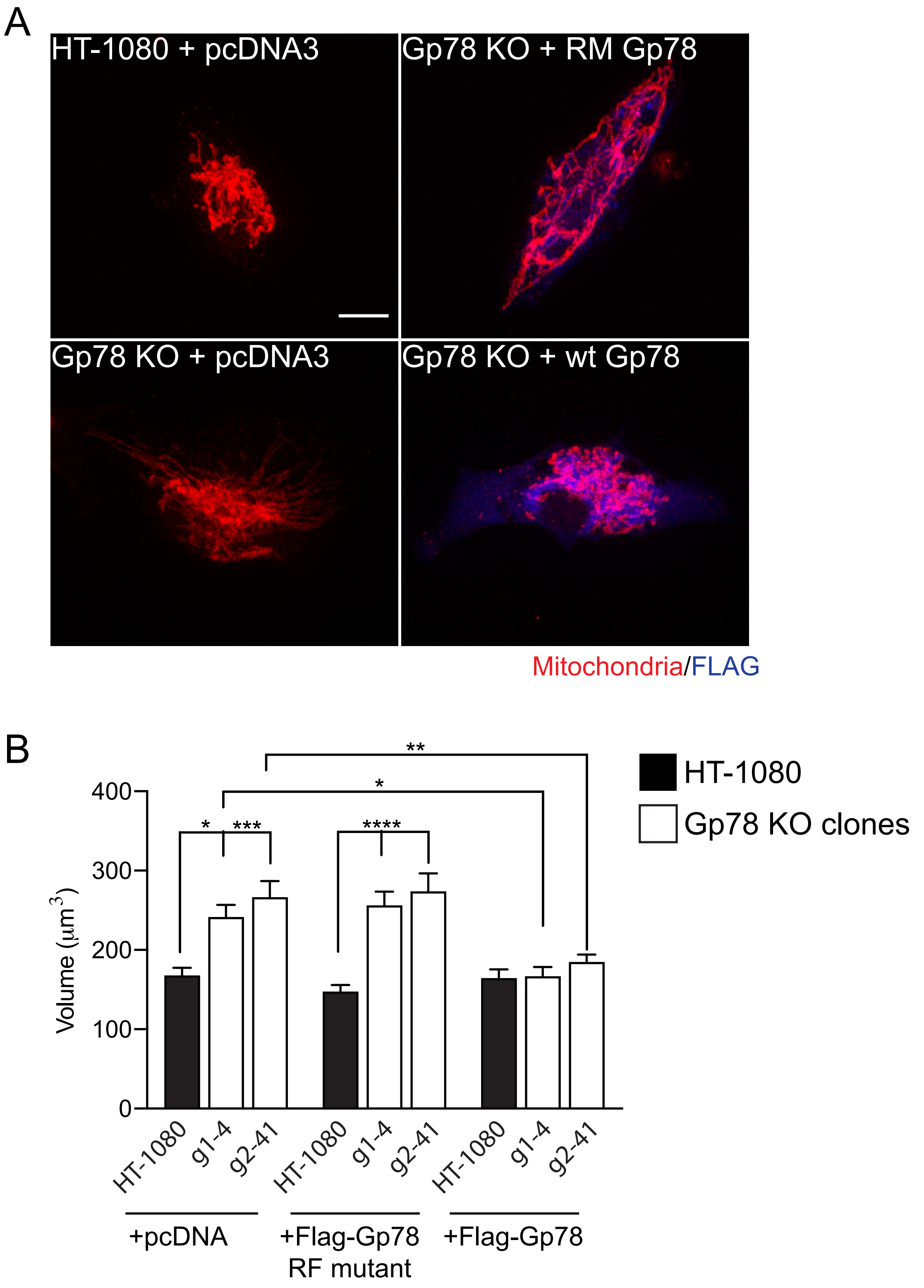


Supplemental Figure 1

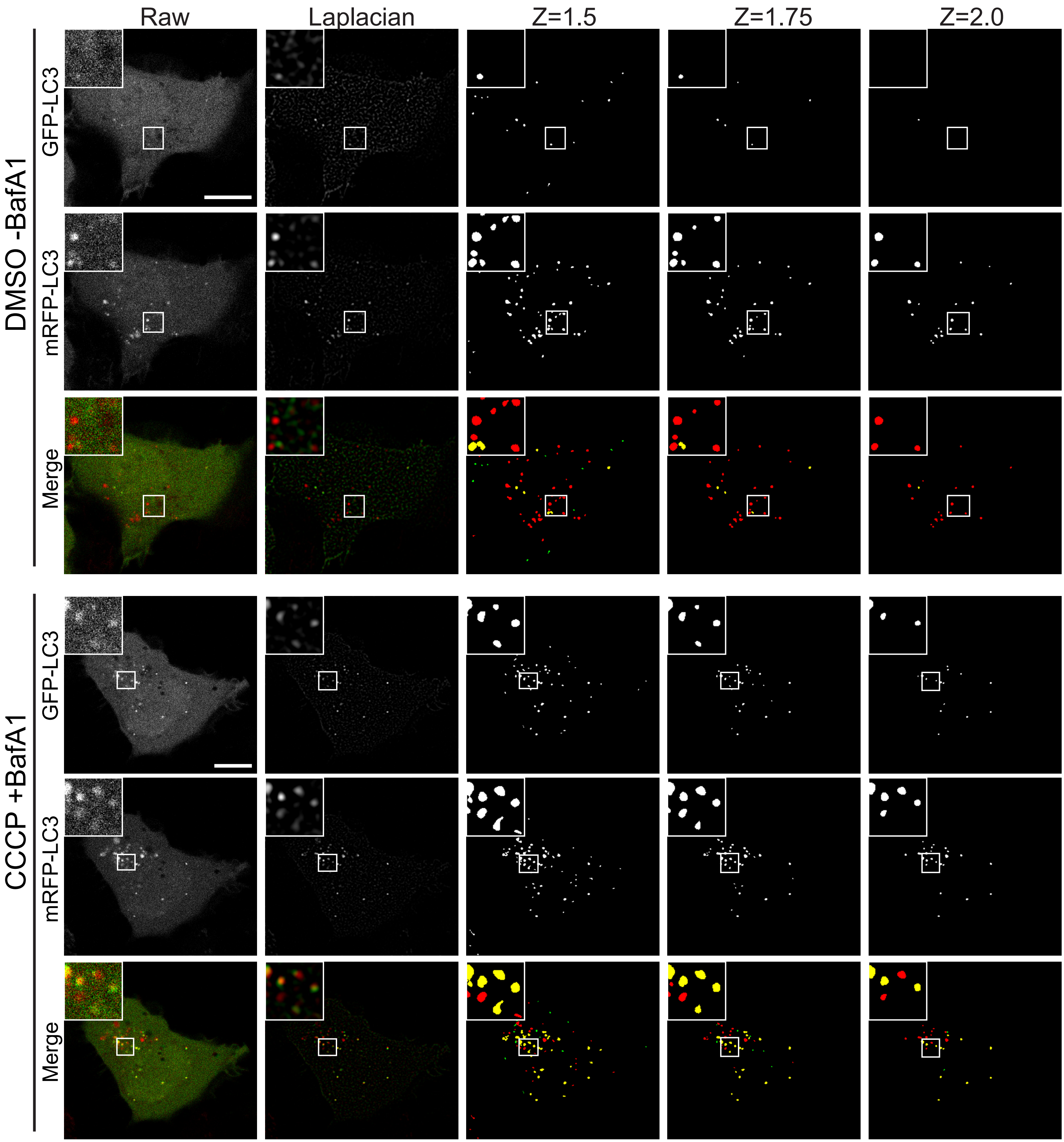


Supplementary Figure 2

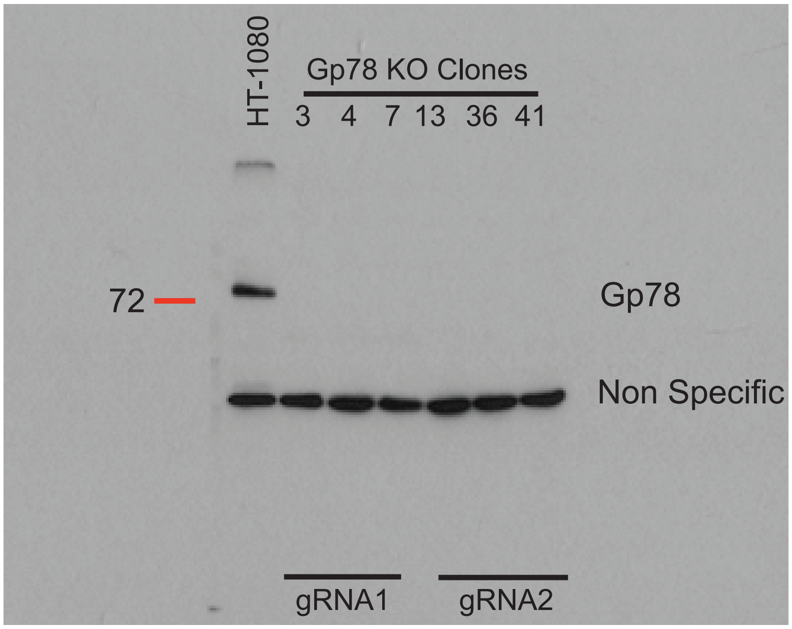


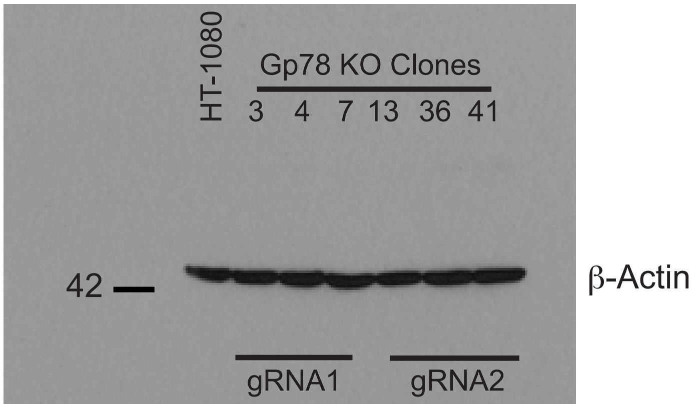


**Supplemental Figure 3 (Full blots for Figure 1A)**

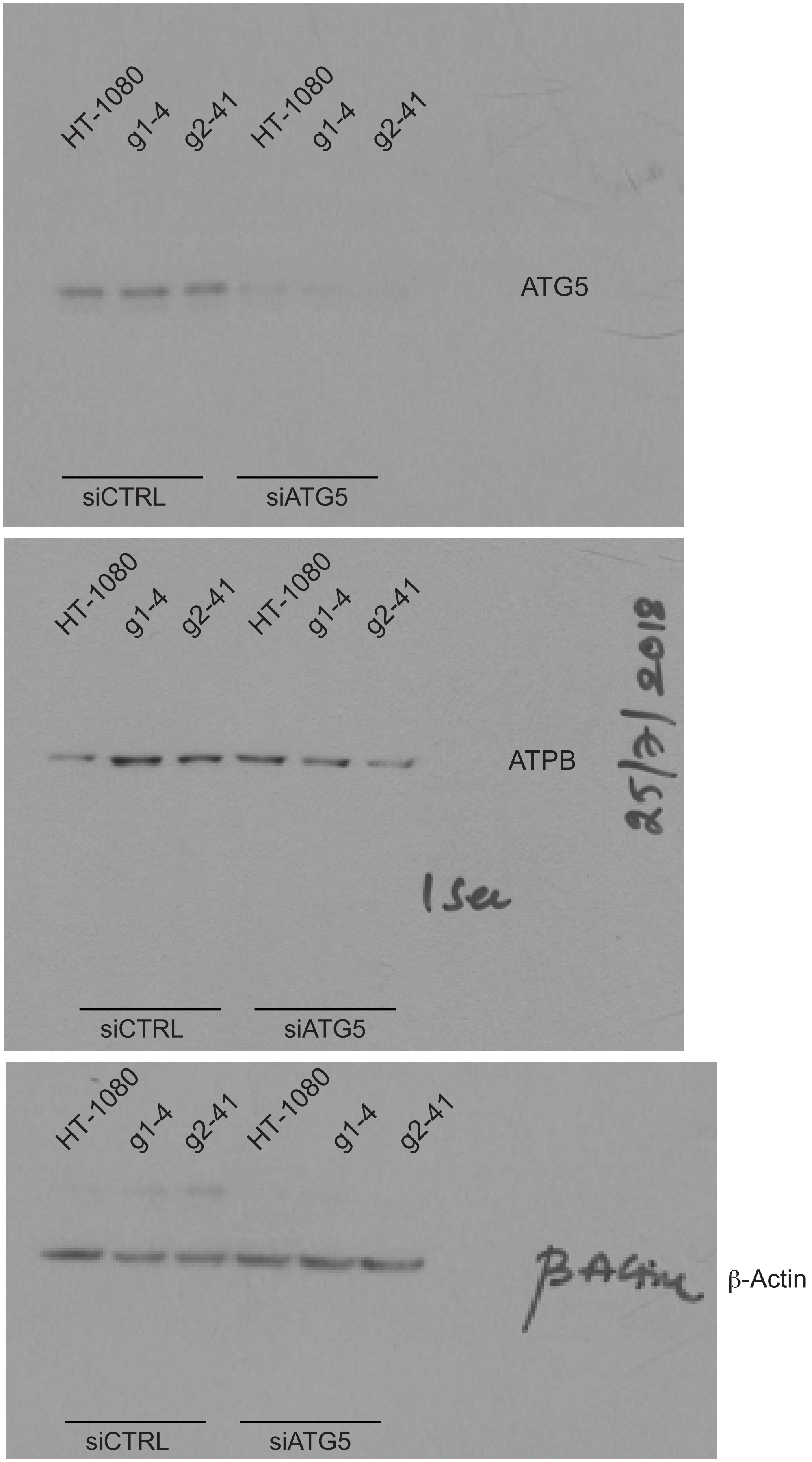


**Supplemental Figure 4 (Full blots for Figure 2A)**

### FIGURE 3-DMSO-CCCP

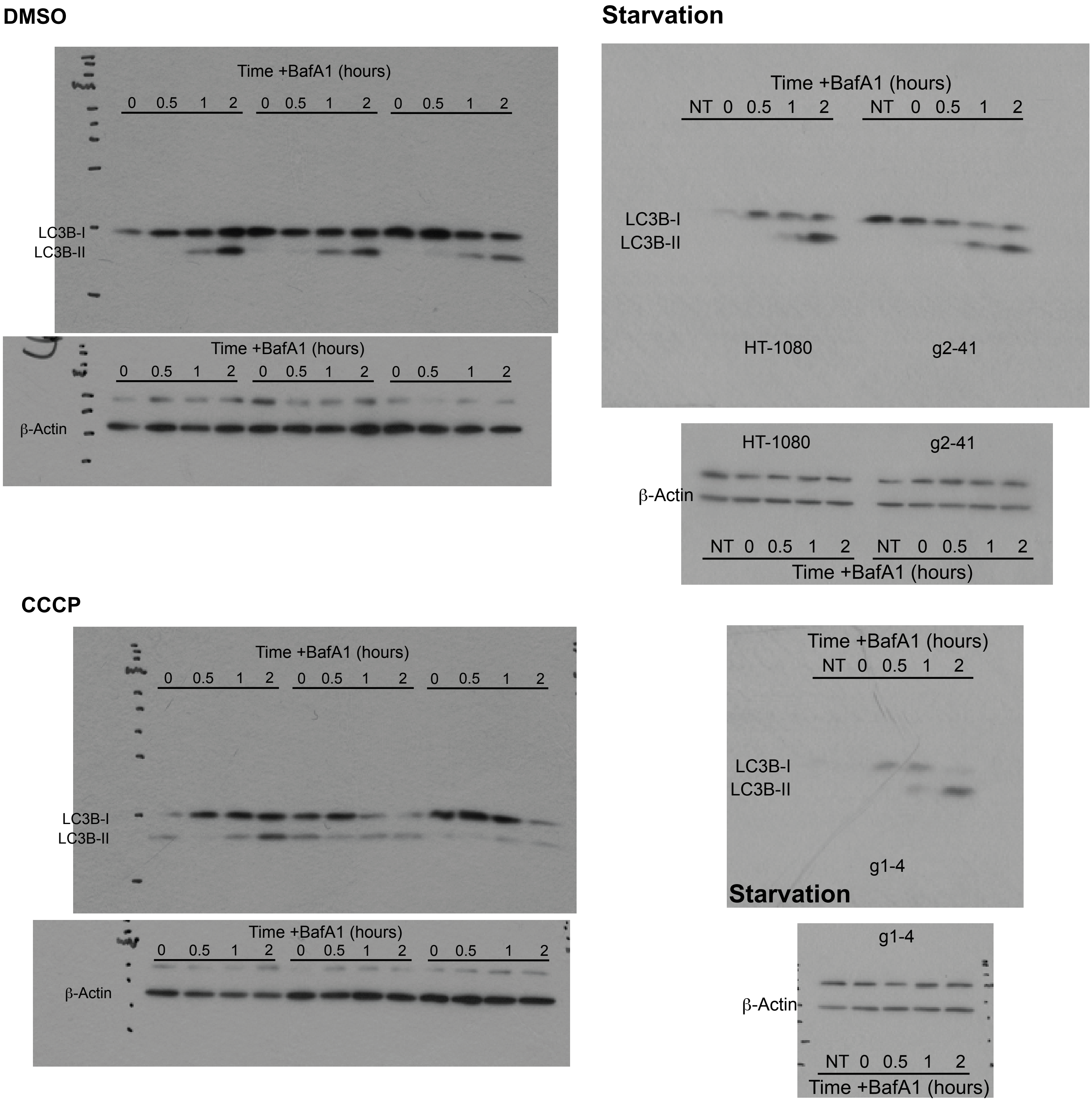


**Supplemental Figure 5 (Full blots for Figure 3)**
